# Network analysis of the papaya orchard virome from two agroecological regions of Chiapas, Mexico

**DOI:** 10.1101/708479

**Authors:** Ricardo I. Alcalá-Briseño, Kena Casarrubias-Castillo, Diana López-Ley, Karen A. Garrett, Laura Silva-Rosales

## Abstract

The study of complex ecological interactions - such as those among host, pathogen, and vector communities - can help to explain host ranges and the emergence of novel pathogens. The analysis of community structures using bipartite networks describe the associations between two trophic levels, for example plants and pollinators, or hosts and parasitoids. Bipartite networks represent interactions (links) occurring only between nodes in different levels - in our case, between viruses and hosts. We evaluated the viromes of papaya orchards (papaya, weeds, and insects) from intensive production of papaya in the Pacific Coastal Plain and the Central Depression of Chiapas, Mexico. Samples of papaya cultivar Maradol, which, like most cultivars, is susceptible to papaya ringspot virus (PRSV), were categorized by symptoms by local farmers (papaya ringspot symptoms, non-PRSV symptoms, or no symptoms). These analyses revealed the presence of 61 viruses, where only four species were shared among both physiographic regions. Nearly 52 complete viral genome sequences were recovered, of which 16 showed homology to known viruses, and 36 shared similarities with different genera including *Potyvirus*, *Comovirus*, and *Tombusvirus* (RNA viruses), and *Begomovirus* and *Mastrevirus* (DNA viruses). We analyzed the network of associations between viruses and host-location combinations, and described ecological properties of the network, such as asymmetry in interactions and nestedness compared to null models. Understanding the network structure informs management strategies, and advances understanding of interactions of hosts and viruses in the agroecological landscape.

**Importance:** Virus-virus interactions in plants can modify host symptoms. As a result, disease management strategies may be unsuccessful if they are based solely on visual assessment and diagnostic assays for known individual viruses. Papaya ringspot virus is an important limiting factor for papaya production, and likely has interactions with other viruses that are not yet known. Using high-throughput sequencing, we recovered known and novel RNA and DNA viruses from papaya orchards in Chiapas, Mexico, categorized by host and, in the case of papaya, symptom type: asymptomatic papaya, papaya with ringspot virus symptoms, papaya with non-ringspot symptoms, weeds, and insects. Using network analysis, we demonstrated virus associations within and among host types, and described the ecological community patterns. Recovery of viruses from weeds and asymptomatic papaya suggests the need for additional management attention. These analyses contribute to the understanding of the community structure of viruses in the agroecological landscape.

## Introduction

Pathogen emergence results from interactions between susceptible hosts and pathogenic viruses in conducive environments, causing disease outbreaks in new geographic regions or hosts (1, 2). Emerging diseases caused by pathogen expansion to new hosts, or pathogen host jumping, are regularly reported in new hosts, vectors and regions, causing yield losses in many parts of the world (3). An example is the emergence of diseases like maize lethal necrosis in sub-Saharan Africa, caused by a synergistic interaction between two single stranded (ss) RNA viruses, a potyvirus and a tombusvirus (4, 5). Another important example is the viral complex of several species of ssDNA begomoviruses that causes cassava mosaic disease, in a pandemic spreading through Africa via whiteflies (6, 7).

A recent disease outbreak of a mixed infection of RNA and DNA viruses, the interaction of potyvirus, crinivirus and begomovirus, caused severe symptoms in papaya (*Carica papaya* L.) orchards in Texas (8). Papaya ringspot virus (PRSV), a plant ssRNA potyvirus vectored by aphid species in a non-persistent manner, or mechanically transmitted through farm tools, causes major yield losses due to foliar deformation, resulting in reduced photosynthetic area, and lowers fruit quality by producing ringspots (9). Previous studies of papaya production areas across Mexico found six different strains of PRSV, and mixed infections with papaya mosaic virus (PapMV) (10, 11). In the 1990s a rhabdovirus was identified in southeast Mexico with similar etiology to papaya apical necrosis disease or papaya droopy fruit (12–15). A disease reported since the 1980s in Brazil, named *meleira disease*, or ‘sticky disease’, of papaya, caused by the papaya meleira virus (PMeV) (16), was reported in Yucatan, Mexico (17, 18). Recently, papaya meleira virus 2 (PMeV-2), with etiology similar to meleira disease (19), was discovered, but is not yet known to cause problems for papaya production in Mexico. Plant-virus interactions can have devastating outcomes, however interactions between viruses can also be antagonistic, as we recently reported in a time series study of co-infection of PapMV and PRSV (20).

Interactions among viruses, including synergistic and antagonistic interactions, may be a common feature in nature (21). Co-infections modify symptoms in different ways, resulting in mild to severe symptoms, or asymptomatic infected plants. Thus, co-infections can produce misleading diagnoses when planting material from tissue culture is tested. Plant-virus interactions can pass unnoticed, and may be difficult to interpret by standard methods such as visual evaluation of symptoms, ELISA, PCR and/or RT-PCR (22). Due to improvement of protocols for nucleic acid isolation of virus-like particles, and availability of high-throughput sequencing technologies, it is now possible to explore all or almost all viruses associated with a given host. Viral metagenomics approaches support the discovery of a large number of virus species in wild plants, crops, and vectors (22–25). In addition to analyses of diversity indices, methods for evaluating associations among pathogens are needed to understand the potentially complex network of interactions among microbes, epidemiological dynamics, and bipartite networks specific to host-vector and plant-virus interactions (26, 27). Network analysis provides insights into biological relationships in viral communities, and can generate hypotheses about mechanisms that promote and prevent the co-occurrence of viruses in communities (28). Systems that include multiple hosts, and potential unknown interactions between RNA and DNA viruses, add another layer of complexity. Bipartite networks have been evaluated extensively for plant-pollinator interactions, which are interesting analogs for host-pathogen systems (29, 30).

Bipartite network analysis can be used to evaluate community nestedness and modularity. In more nested communities, the viruses associated with one host will tend to be a subset of the viruses associated with another host. In modular communities, nodes tend to be divided into subsets, forming modules, such that one group of viruses tends to infect one group of hosts. Theoretical studies have shown that a nested network structure can minimize competition, increasing the number of coexisting species, and potentially making nested communities more robust to random extinctions (31). The network of specialization has characteristics, such as modularity or compartmentalization patterns, that supports the network stability between generalist and specialist species (31).

Our analysis synthesizes two conceptual frameworks: viral metagenomics to reveal plant viruses using high-throughput sequencing, combined with analysis of plant viruses in bipartite networks. Using concepts from island biogeography (32), we defined an environment as a combination of host type, in the broad sense, and region. This analysis shows how hosts connect virus species through the network, and how plant–vector-virus associations can be interpreted to inform management strategies for papaya orchards in Chiapas.

In the papaya orchard virome network, the link between PRSV and papaya plants displaying typical symptoms of ringspot is expected in both regions. Also, it is expected that some viruses will be nested within papayas and weeds, or papayas and insects, indicting potential management strategies. However, what other types of interactions are present in the papaya orchard virome? We used metagenomics to characterize the papaya orchard virome, and network analysis to characterize viral associations across the host types in two physiographic regions in southeast Mexico (Fig. 1), separated by the Sierra Madre de Chiapas mountain range at 4,093 m elevation (33). These two physiographic regions represent different ecosystems, sharing some characteristics but differing in the degree of land fragmentation, orchard size, and plant species, among other characteristics (34). The first, the Central Depression, is a tropical deciduous forest at 700 m above sea level. The second, the Pacific Coastal Plain, is a highly fragmented semi-deciduous tropical forest, at sea level, with forests being replaced to a great extent by grasslands for cattle grazing. Both physiographic regions are located in the Federal State of Chiapas, one of the main producers and exporters of papaya in southern Mexico (<u>INAFED</u>). Management strategies for papaya orchards in these regions emphasize reducing PRSV effects, and roguing of infected papaya is the primary strategy to control ringspot disease.

**Fig 1.**
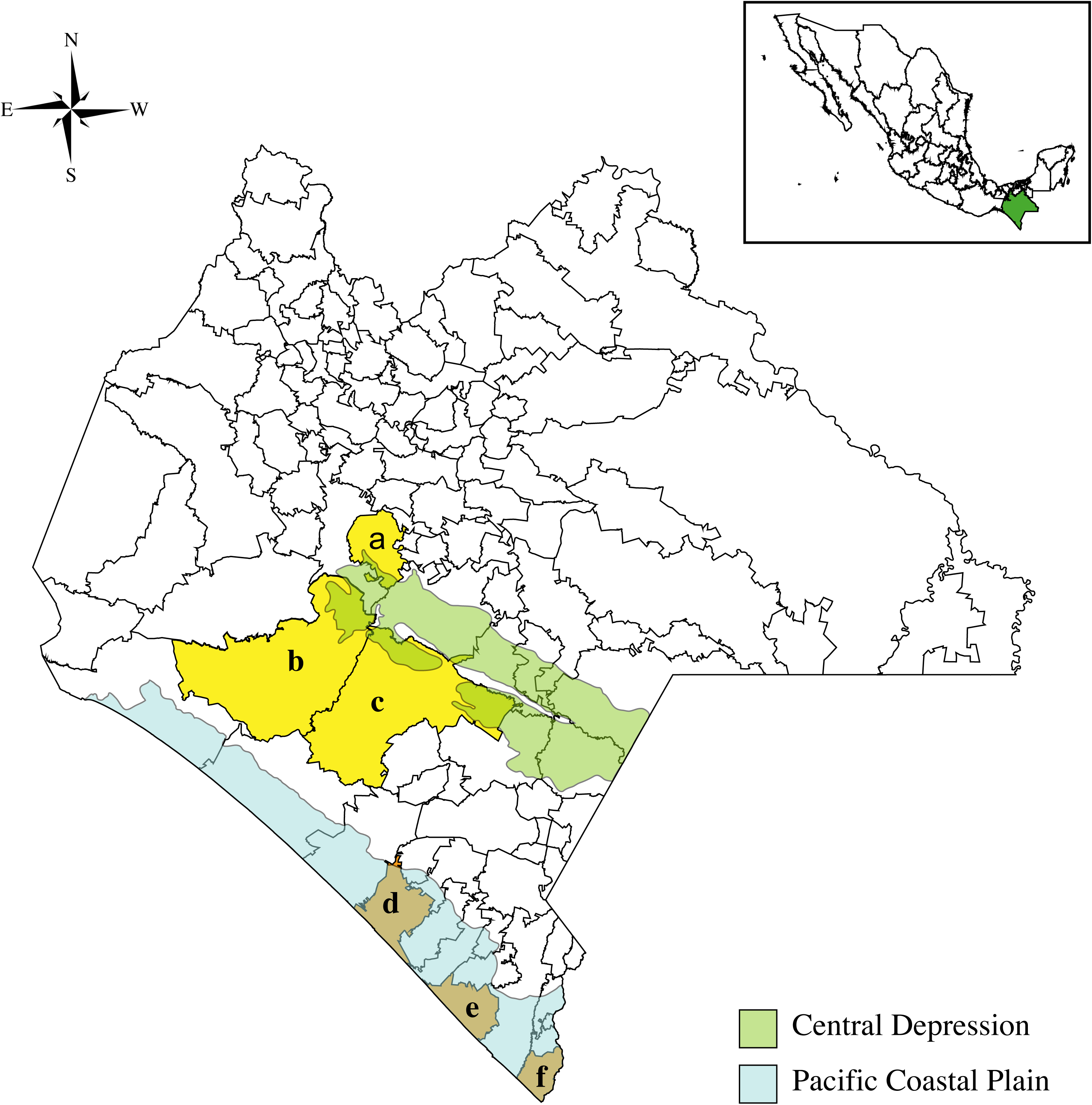
Two physiographic regions in the Federal State of Chiapas in Mexico. Central Depression (at 700 masl; light green), and Pacific Coastal Plain (at sea level; light blue) and the counties from which samples were collected: a) Acalá, b) Villa Corzo, c) La Concordia, d) Acapetahua, e) Mazatán, and f) Suchiate.

Here we (i) report 52 near full-length genome sequences identified as plant viruses, 29 being novel viruses, associated with papaya, weeds, and insects, (ii) report the diversity and distribution of viruses in two contrasting physiographic regions, the Central Depression and the Pacific Coastal Plain of Chiapas, and (iii) evaluate the bipartite network structure of viruses and host-location combinations in the papaya orchard agroecosystem. The papaya orchard virome network has structures beyond stochastic associations. Bipartite network analysis of these complex viromes, generated through viral metagenomics, revealed virus associations and identified the roles of particular hosts in connecting the network.

## Results

To evaluate the papaya orchard virome, papaya samples were collected based on symptoms evaluated by local growers. Growers are generally familiar with PRSV symptoms because management in orchards commonly consists of roguing papaya plants with typical PRSV symptoms or other virus symptoms. To evaluate where PRSV and other viruses were present in papaya orchards, we used a viral metagenomics approach to assess the diversity of viruses in the region. We identified 82 sequences corresponding to at least 57 viruses, of which 52 sequences were nearly full viral genomes within 10 genera, and 10 were unassigned genera in 10 viral families. The bipartite network of viruses, based on metagenomic results, and host-location combinations for the papaya orchards, was evaluated in terms of modularity and nestedness, and centrality measures (Box 1).

### Box 1. Metric definitions

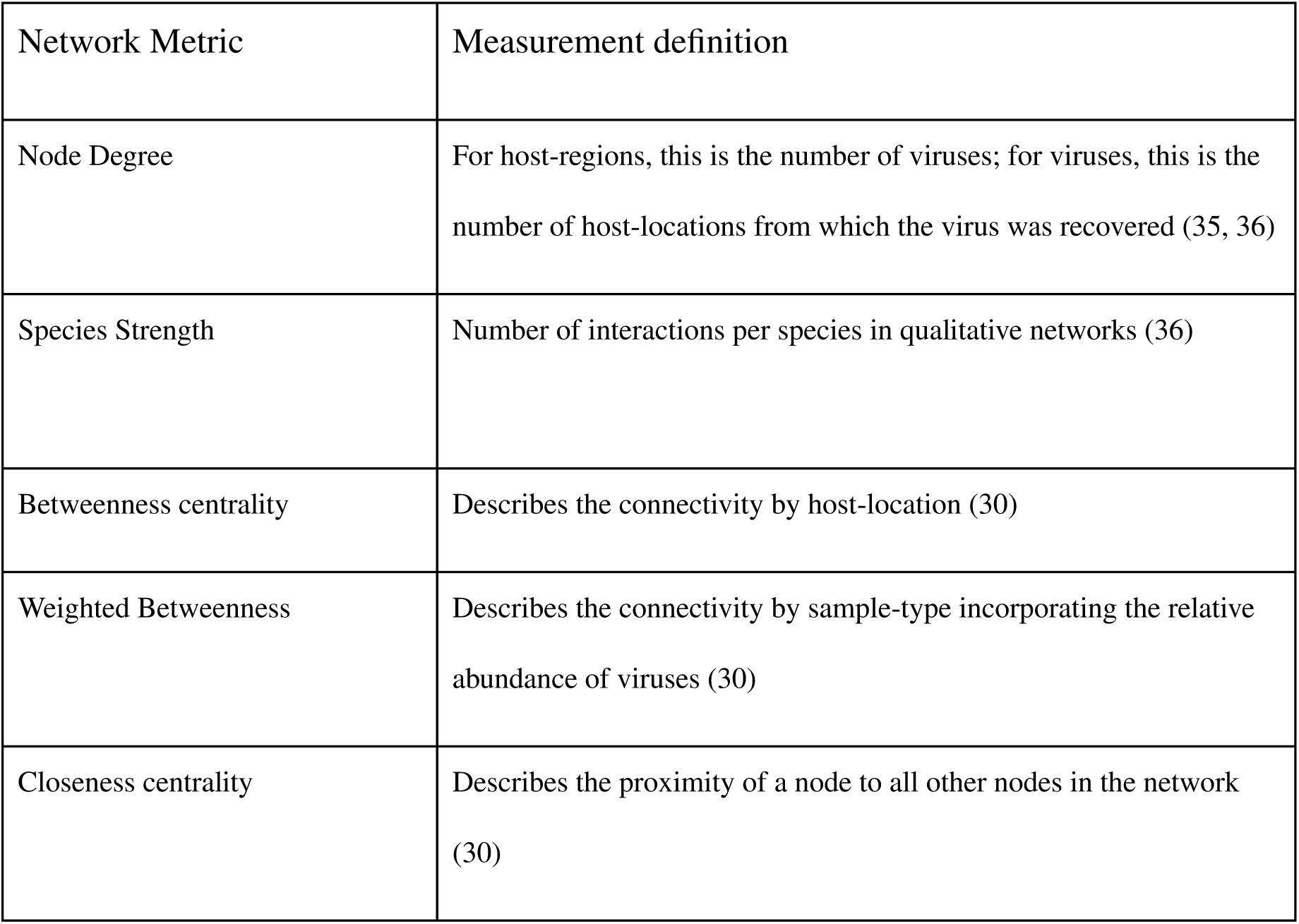
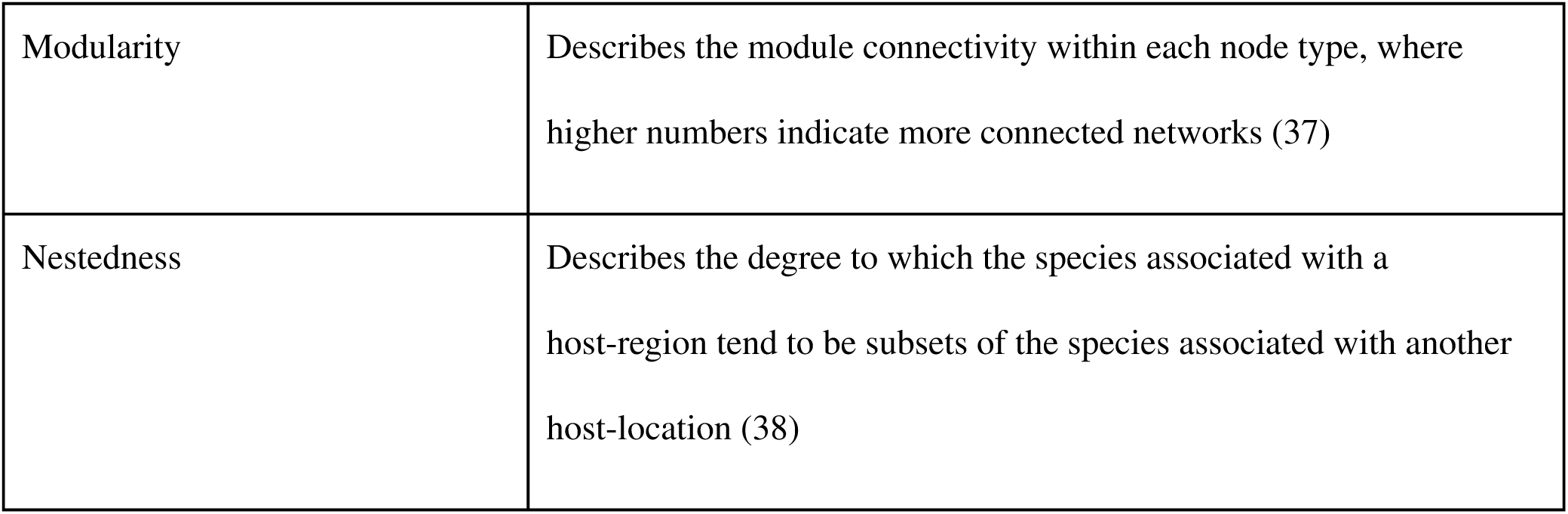

Bipartite networks have frequently been described for cases such as pollinator networks, where the two types of nodes are pollinators and plants (29). We conceptualize bipartite networks for the papaya orchard virome as having one level representing hosts (in the broad sense: the primary host (papaya, divided into three categories based on symptoms), secondary hosts (weeds), and vectors (insects), for each of two geographic regions, and a second level representing viruses. A link between a virus and a host-location combination indicates that the virus was present in that host in that location. These analyses clarify the interactions among RNA and DNA viruses in papaya production areas, and the prevalence and distribution of viruses in secondary hosts and insect vectors. Understanding these relationships can inform strategies for management of viruses that pose a risk to papaya production, and can contribute to assessments of the risk of virus spillovers.

### Viral-like particles

The diversity of viruses was first evaluated by observing the purified viral-like particles (VLPs) with an electron microscope. Several viral particle morphotypes were confirmed, including filamentous, icosahedral and pleomorphic. We estimated the diameters for icosahedral particles as ranging from 15 to 65 nm and lengths of up to 750 nm for filamentous particles. Viral morphologies resembling flexuous filamentous or rigid helical rods particles were observed in papaya plants from both regions, the Pacific Coastal Plain and Central Depression. A large number of icosahedral particles were observed in all samples. The presence of pleomorphic particles suggested the presence of rhabdoviruses in papaya plants from the Pacific Coastal Plain (Fig. S1).

### Overall composition and relative abundance of plant viruses present in papaya orchards

There were 82 sequences with homology to viruses, and 61 unique viral sequences of 56 viruses. Only three virus species (four viral sequences) were recovered from both physiographic regions: PRSV, PapMV and Euphorbia mosaic virus (EuMV). Thirty-three virus species (53% of the total number of virus-location-host type combinations) were obtained from samples from the Pacific Coastal Plain, and 32 viruses (47%) from the Central Depression (Fig. 2). The percentage of DNA and RNA viruses was calculated using the presence and sequence coverage as virus relative abundance (Fig. 2). The percentage of virus-location-host type combinations for asymptomatic papaya was similar in the Pacific Coastal Plain (20.6%) and the Central Depression (19.3%). However, samples from papaya with other viral symptoms showed differences in virus-location-host type combination percentages, 12.9% and 2.8% respectively. Papaya identified by growers as having symptoms of PRSV was presenting similar percentages, 19.5% in the Pacific Coastal Plain and 24.9% in the Central Depression. Interestingly, only samples from papaya with symptoms of PRSV were associated with both RNA and DNA viruses in both regions; samples from asymptomatic papaya and papaya with other type of symptoms were associated only with RNA viruses (Fig. 2a). The plant virus-location-host type combinations associated with weed samples (including *Euphorbia* sp.*, Ipomea* sp.*, Sida* sp.*, Portulaca* sp., wild grasses, and maize; for a list of weeds, see Table S1), were 9.4% RNA viruses and 27.7% DNA viruses for the Pacific Coastal Plain, and 17.9% RNA viruses and 44.9% DNA viruses for the Central Depression (Fig. 2b). The plant virus-location-host type combinations associated with the insect samples (including insect orders such as Coleoptera, Hemiptera, and Dermaptera; Table S2), were 69.4% RNA viruses and 8.3% DNA viruses for the Pacific Coastal Plain, and 20.3% RNA viruses and 1.97% DNA viruses for the Central Depression (Fig. 2c).

**Fig 2.**
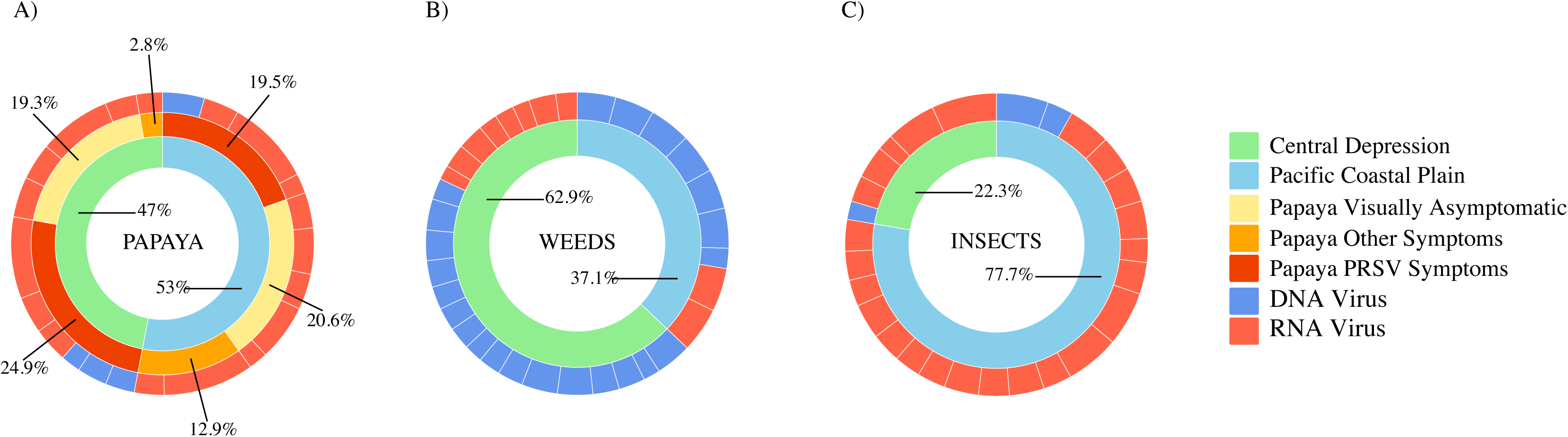
Distribution of DNA and RNA viruses in papaya, weeds and insects. The frequency of log-transformed relative sequence abundance of viruses from the papaya orchard virome by host-location: A) papaya plants, B) weeds and C) insects. The inner ring indicates the proportion of viruses from the samples collected from the Pacific Coastal Plain (light green) and the Central Depression (light blue). The middle ring (papaya only) indicates the proportion of viruses for three different types of papaya symptoms: visually asymptomatic (yellow), non-PRSV symptoms (orange), and PRSV symptoms (red). The outer ring indicates the proportion of putative RNA (light red) and DNA (blue) virus species.

### Virus classification

All viruses were grouped taxonomically within ten families, with ten approved genera and twelve viruses that could not be classified at genus level. Nine viruses were identified from papaya, in six genera: *Comovirus*, *Crinivirus*, *Potexvirus*, *Potyvirus*, *Nucleorhabdovirus*, and *Begomovirus*, and an unclassified dsRNA virus, PMeV-2, a toti-like virus. In the Pacific Coastal Plain, potexviruses, comoviruses, potyviruses, nucleorhabdoviruses and begomoviruses were present, and potexviruses, criniviruses, potyviruses, unclassified dsRNA viruses and begomoviruses were present in the Central Depression (Fig. 3a, bottom bars). In the Pacific Coastal Plain, asymptomatic papaya samples included potexviruses, comoviruses, potyviruses, and nucleorhabdoviruses; samples from papaya with non-PRSV symptoms yielded sequences identified as potyviruses and nucleorhabdoviruses; and samples from papaya with PRSV symptoms sequences were identified as potexviruses, potyviruses, nucleorhabdoviruses, and begomoviruses (Fig. 3b, upper bars). In the Central Depression, asymptomatic papaya yielded sequences identified as criniviruses, potyviruses and PMeV-2; papaya with non-PRSV symptoms yielded one virus identified as PapMV, a potexvirus; and papaya with symptoms of PRSV yielded sequences identified as criniviruses, potyviruses, and begomoviruses (Fig. 3b, bottom bars). Thirty-one viruses were recovered from weeds in both regions and classified in four RNA genera (*Comovirus*, *Potyvirus*, *Potexvirus*, and *Waikavirus)*, in two DNA genera (*Begomovirus*, and *Mastrevirus*), as two novel unclassified potyviruses, as one unclassified caulimovirus, as a DNA pararetrovirus, and a satellite virus (Fig. 3a, upper bars). The alphasatellite sequence (Acc. No: pending) showed 78% similarity with the dragonfly-associated alphasatellite characterized previously in Puerto Rico (39). Twenty-seven sequences were recovered from insects, four of them near-full length genomes, and classified in five RNA genera: *Comovirus*, *Potyvirus*, *Tombusvirus*, *Tymovirus*, *Waikavirus*. Two DNA genera were recovered from insects, *Begomovirus* and *Mastrevirus*, along with unclassified alphaflexiviruses and tymoviruses (Fig 3a, middle bars).

**Fig 3.**
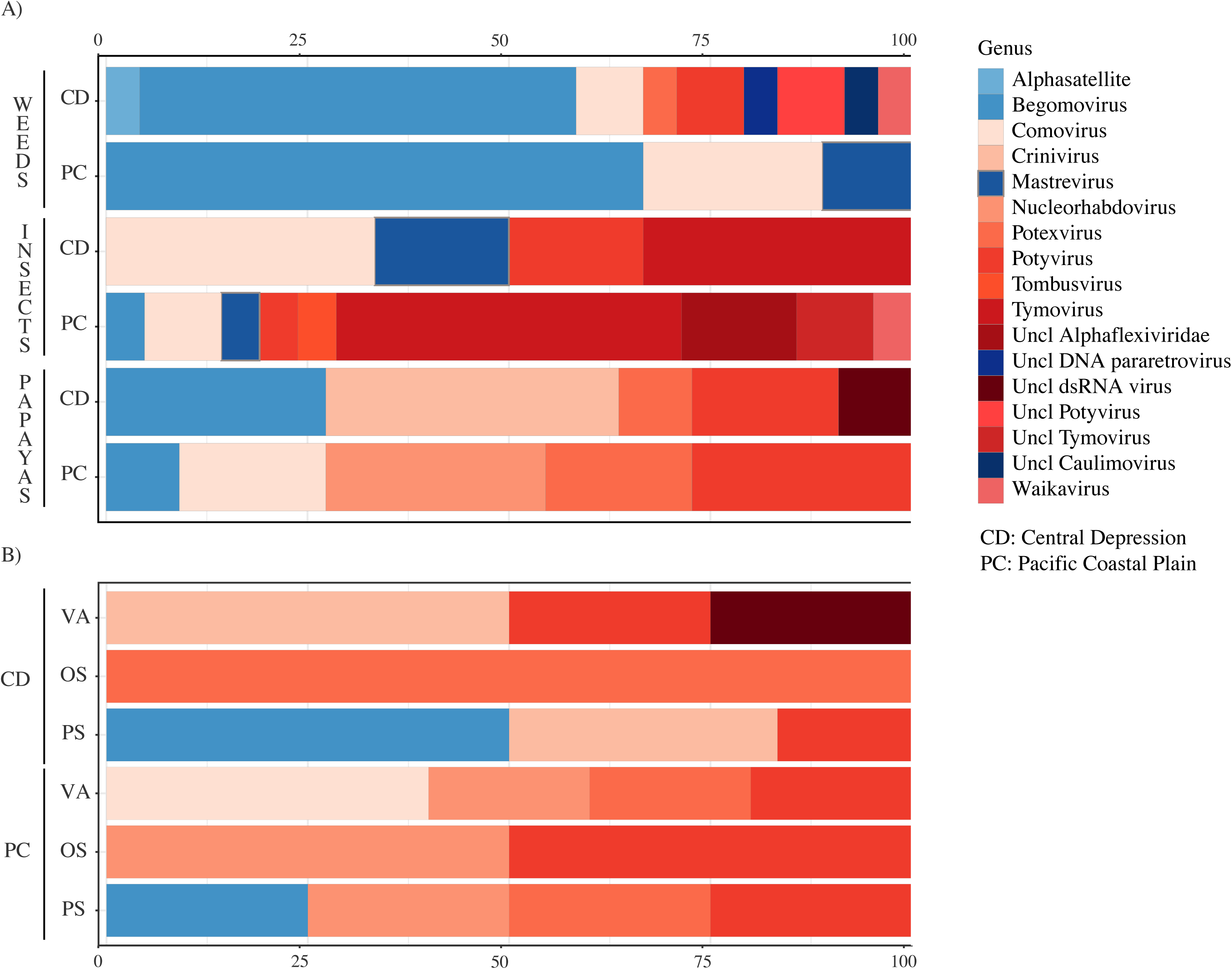
Relative abundance of DNA and RNA by genus. The log-transformed relative abundance of DNA and RNA viruses by genus and by host-location in the papaya orchard virome of Chiapas. Panel A: the diversity and relative abundance of viruses by host-location for weeds, insects and papaya plants, by physiographic region for the Central Depression (CD) and Pacific Coastal Plain (PC). Panel B: the diversity and relative abundance of viruses in papaya divided by symptoms: visually asymptomatic (VA), non-PRSV symptoms (OS), and PRSV symptoms (PS). Cold colors represent DNA viruses, and warm colors indicates RNA viruses.

### Analysis of virome networks in papaya orchards

The papaya orchard virome network included 61 viral sequences and 82 links connecting virus nodes and host-location combination nodes (Fig. 4). The number of links (node degree) for a region, equal to the number of viruses recovered from samples from that region, was 33 for the Pacific Coastal Plain, and 32 for the Central Depression. Only four (viruses) nodes were shared by both regions: PRSV, PapMV, EuMV-A, and EuMV-B.

**Fig 4.**
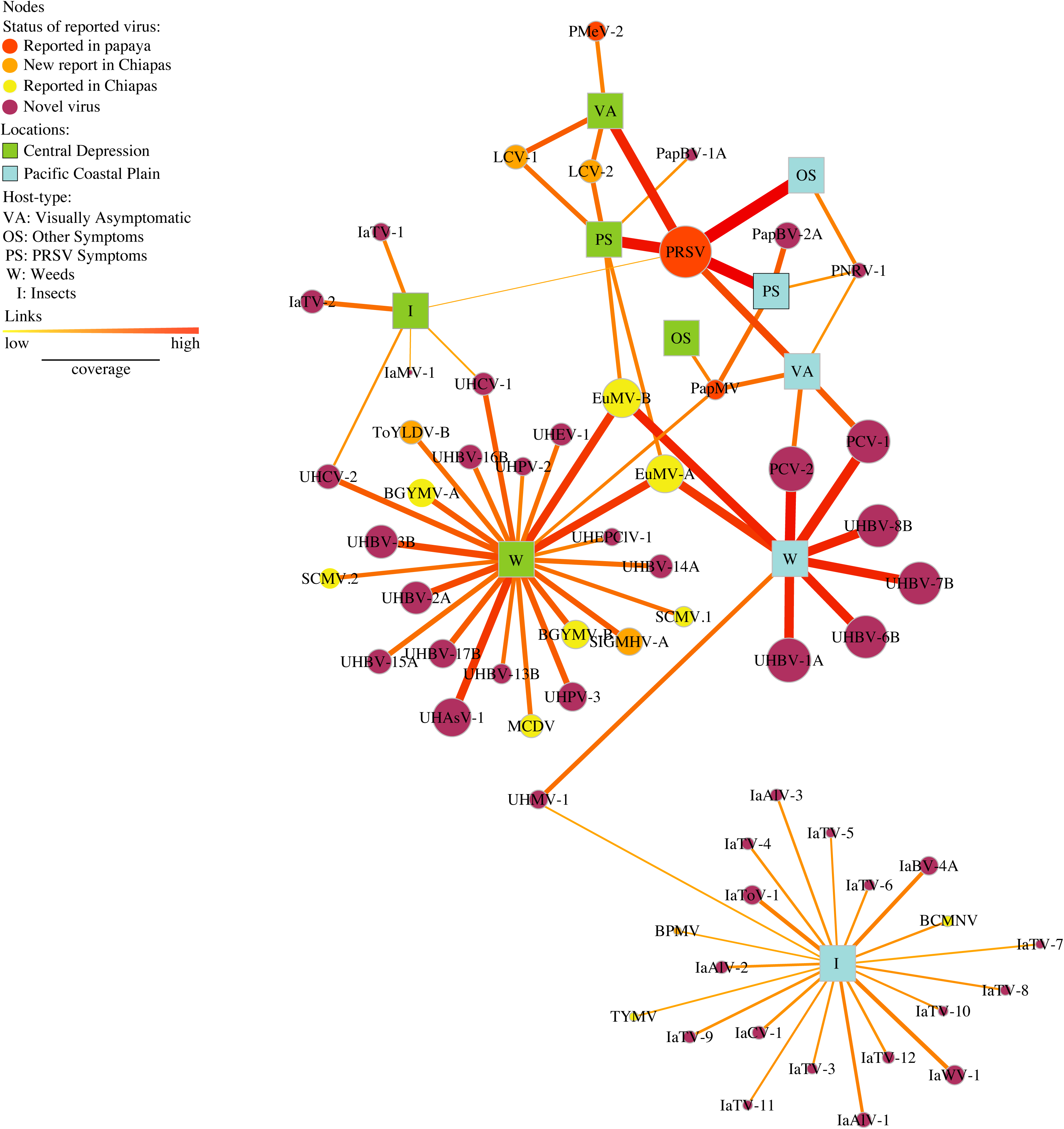
Bipartite network analysis of the papaya orchard virome. Papaya orchard virome, represented in a bipartite network with two types of nodes: viruses (circles) and host-location combinations (squares) for two physiographic regions, the Pacific Coastal Plain (light blue) and Central Depression (light green). The size of circular nodes varies with the sum of the relative abundance by virus specie. Links indicate that a virus is present in a host-location, and their width and color (yellow to red) represent relative abundance by host-location. Alternate hosts are weeds (W), insects (I), papaya with PRSV symptoms (PS), papaya with non-PRSV (OS), and visually asymptomatic (VA). Viruses are indicated by their acronyms. Node color represents the status of the virus: reported in papaya (red), new report for Chiapas in non-papaya hosts (orange), previously reported in Chiapas in non-papaya hosts (yellow); and novel viruses not yet reported anywhere else (purple).

The node degree for each host-location combination indicates the number of viruses recovered from each group (Table 1, Fig. 4). The node species strength is a weighted version of the node degree, the weighted average of the number of interactions of virus by host (36). The highest node strength in the Central Depression was observed for weeds, and in the Pacific Coastal Plain for insects. The lowest node strength observed for the Pacific Coastal Plain was for asymptomatic papaya, and in the Central Depression for papayas with non-PRSV symptoms (Table 1).

**Table 1.** Diversity and network indices.

| Host<br>type | Network metrics |  |  |  |  |  |
| --- | --- | --- | --- | --- | --- | --- |
|  | degree | species<br>strength | betweenness | weighted<br>betweenness | closeness | closeness<br>weighted |
| Pacific Coastal Plain |  |  |  |  |  |  |
| VA | 5 | 0.586 | 0.270 | 0.181 | 0.122 | 0.001 |
| OS | 2 | 1.193 | 0.000 | 0.000 | 0.098 | 0.001 |
| PS | 4 | 1.840 | 0.090 | 0.168 | 0.112 | 0.001 |
| W | 9 | 8.259 | 0.333 | 0.272 | 0.093 | 0.001 |
| I | 21 | 20.058 | 0.000 | 0.000 | 0.060 | 0.000 |
| Central Depression |  |  |  |  |  |  |
| VA | 4 | 2.212 | 0.000 | 0.000 | 0.098 | 0.001 |
| OS | 1 | 0.114 | 0.000 | 0.000 | 0.084 | 0.000 |
| PS | 6 | 2.002 | 0.159 | 0.246 | 0.115 | 0.001 |
| W | 24 | 21.674 | 0.125 | 0.129 | 0.108 | 0.001 |
| I | 6 | 3.057 | 0.020 | 0.000 | 0.105 | 0.000 |

Betweenness centrality indicates the importance of a species as a connector forming bridges between hosts. We calculated an unweighted and weighted version of betweenness centrality. The highest betweenness centrality (unweighted and weighted) was observed for papaya with PRSV symptoms (0.090 and 0.168), visually asymptomatic papaya (0.270 and 0.181), and weeds (0.333 and 0.272) in the Pacific Coastal Plain, and papaya with PRSV symptoms (0.159 and 0.246) and weeds (0.125 and 0.108) in the Central Depression. The rest of the nodes had no role in connecting the network (Table 1). To compare the weighted and unweighted betweenness values, we computed Kendall’s Tau (τ) correlation coefficient (τ= 5.8, corr= 0.9, p= 0.0004), indicating limited differences between the ranks for unweighted and weighted versions of betweenness centrality.

Closeness centrality is a measure of the proximity of one node (host-location) to all other nodes (host-locations) in the network. Surprisingly, asymptomatic papaya in the Pacific Coastal Plain had the highest closeness centrality (0.122) in the network, followed by papaya with PRSV symptoms (0.112 and 0.115) for the Pacific Coastal Plain, and Central Depression, however the weighted closeness was significantly lower (0.001) (Table 1). For the Kendall’s Tau (τ) correlation coefficient, there was some evidence (τ= 2, corr= 0.58, p= 0.07) for differences between the unweighted and weighted versions of closeness centrality (Box 1).

### Network analysis and viral metagenomics reveal hidden associations in papaya orchards

Viral metagenomics revealed previously unknown viruses in the papaya orchard, and their associations with other known viruses such as PRSV. Network metrics indicate host associations with viruses. We can show more clearly how host-locations are linked to each other by removing virus nodes that are linked to only a single host-location (Fig. 5). PRSV was present in all papaya sample types in both regions with the exception of non-PRSV symptoms in the Central Depression, where only PapMV was present. PRSV was also associated with insect samples from the Central Depression. On the other hand, PapMV was present in papaya with symptoms of PRSV, and asymptomatic papaya in the Pacific Coastal Plain, as well as in weeds in the Central Depression. A putative papaya nucleorhabdovirus 1 (PNRV-1) was present in all samples of papaya in the Pacific Coastal Plain, and associated with PapMV and PRSV. Novel putative bipartite comoviruses, papaya comovirus (PCV) 1 and 2, were identified in weeds and were also present in asymptomatic papaya in the Pacific Coastal Plain. A novel mastrevirus provisionally named unknown host mastrevirus 1 was isolated from weeds and insects in the Pacific Coastal Plain. EuMV (*Begomovirus*) was identified in weeds from both regions, but only in the Central Depression was it associated with PRSV and Lettuce chlorosis virus (LCV, *Crinivirus*), in papaya showing symptoms of PRSV. Interestingly, PRSV and LCV were present in asymptomatic papaya in the Central Depression. A novel bipartite comovirus was recovered from both weeds and insects in the Central Depression, and was different from PCV from the Pacific Coastal Plain and two comoviruses tentatively named unknown host comovirus 1 (UHCV-1) and 2 (UHCV-2).

**Fig 5.**
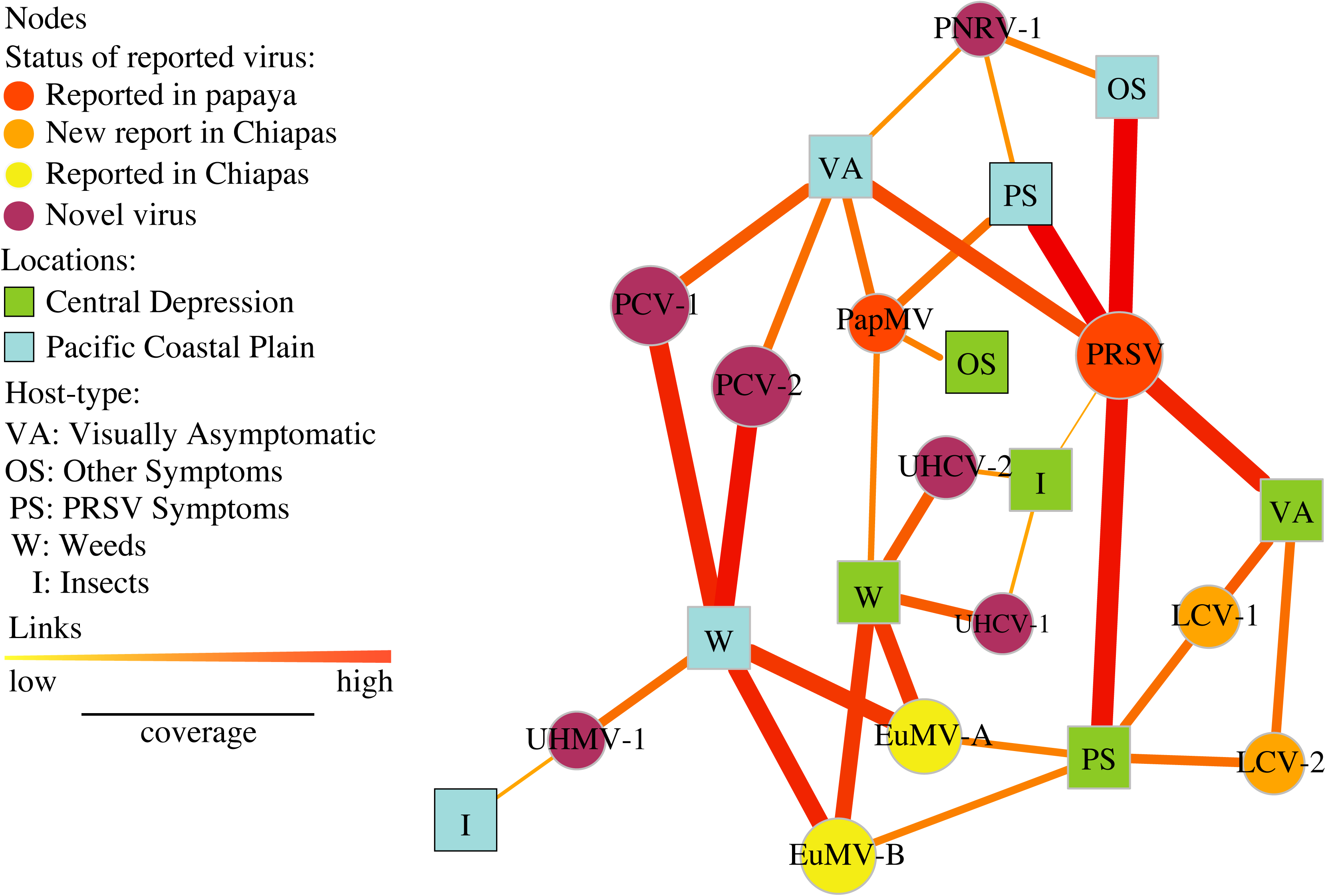
Bipartite network displaying only the nodes that have two or more links. Viruses (circles) and host-location combinations (squares) from two physiographic regions, the Pacific Coastal Plain (light blue) and Central Depression (light green) are represented. Node sizes are proportional to the sum of the relative abundance of virus species. Links indicate associations between nodes, and their width and color (yellow to red) represent low and high relative abundance, respectively, by host-location. Hosts are weeds (W), insects (I), papaya with PRSV symptoms (PS), non-PRSV symptomatic papaya (OS), and visually asymptomatic (VA). Viruses are indicated by their acronyms. Node color represents the status of the virus: reported in papaya (red), new report for Chiapas in non-papaya hosts (orange), previously reported in Chiapas in non-papaya hosts (yellow), and novel viruses not yet reported elsewhere (purple). The PapMV node was added, even though unlinked due to being present only in OS Central Depression.

### Analysis of the virome community structure in papaya orchards

We used two network metrics to summarize the viral community structure: modularity and nestedness (Box 1). These metrics were calculated using the package bipartite (29) in the R programming environment (40). We tested whether there was evidence that the observed virus community patterns, in terms of these two metrics, differed from what would be expected under three null models (41). These three null models were used to generate new adjacency matrices based on specific properties of the observed data. Under null model 1, species richness is maintained for each host-location combination. Reshuffling occurs within host-location combinations, such that the specific viruses associated with each host-location can change while the number of viruses associated with the host-location remains the same. Under null model 2, the number of host-locations in which a species is found is maintained. Reshuffling occurs within each virus species, such that the specific host-locations associated with each virus can change while the number of host-locations for each virus remains the same. Under null model 3, neither species richness nor the number of host-locations associated with a virus are maintained, so reshuffling occurs across both. Comparing results for the three null models can be used to interpret whether deviations from random network structures may be due simply to “first order” properties (such as the total number of species) (29). For example, if null model 1 is rejected but null model 3 is not rejected, this indicates that first order properties alone do not explain the patterns (29). For each null model, the observed adjacency matrices were reshuffled 1000 times, and the observed values were compared to 10,000 random null matrices using a Z-score test.

The modularity of the network describes the connectivity within each node type, where the more interactions there are between host-location nodes, the more connected is the network. The observed modularity for the papaya orchard virome was 0.37, where by comparison modularity values observed in other real networks range between 0.3 and 0.7 (42). There was strong evidence to reject null model 1 (p < 0.001), but little evidence to reject null model 2 (p = 0.216) and null model 3 (p = 0.213) (Table 2). The nestedness of the network indicates the degree to which, for example, the viruses associated with one host-location tend to be a subset of the viruses associated with another host-location. The observed nestedness was 0.576, suggesting a moderate level of nestedness. There was strong evidence for rejecting all three null models for nestedness (Table 2). In general, the low level of modularity and the nestedness of the papaya orchard virome reflect the module formed by papaya plants across symptomatologies, and the nestedness patterns for weeds and papaya plants, and for weeds and insects, in both locations.

**Table 2.** Network metrics.

|  | Observed | Null 1<br>p-value | Null 2<br>p-value | Null 3<br>p-value |
| --- | --- | --- | --- | --- |
| Modularity | 0.373 | $4.32e^{-34}$ † | 0.216 | 0.213 |
| Nestedness | 0.576 | $3.356e^{-07}$ | $4.533e^{-14}$ | 0.0047 |

†not normally distributed

## Discussion

Expanding virome databases offer new opportunities for analysis and understanding of virome complexity. Typical approaches to virome analysis include use of descriptive statistics, sequence analysis for the identification and discovery of novel viruses, and studies of evolution and viral diversity (43–49). More recently, some studies have captured the association of viruses and hosts using network analysis by mining databases or ELISA (50, 51). Here we present a new application of bipartite network analysis coupled with viral metagenomics in a framework that reveals interactions of the entire community, including both known and new virus associations across hosts. Node centrality measures, such as node degree, betweenness, and closeness, provide information to motivate follow-up analyses of management strategies, and more generally provide new insights into ecological interactions between viruses. There was strong evidence that the papaya orchard virome had a nested structure, but only weak evidence for modularity. The patterns of nestedness in this virome indicate subsets of viruses and host-locations that may need to be managed together, where one host-location combination may act as a risk factor for another. To represent this, the bipartite network was converted to a one mode network for the host levels, where modularity and nestedness patterns are connecting the different for host-location combinations emphasizing interactions. For example, we found viruses in asymptomatic papaya in both regions, suggesting that roguing papayas with viral symptoms may not be sufficient to manage virus incidence. We also found PapMV in weeds in the Central Depression, and a novel comovirus (UHCV) emerging in papaya found in weeds in the Pacific Coast, suggest that management strategies could expand to weed control, removing potential hosts for known or emerging pathogens of papaya.

### Many viruses but few interact with papaya

*Carica papaya* is the only species in the family Caricaceae, a cultivated species originating in southern Mexico (52). Papaya viruses tend to be specialists, but some also may infect relatives of *C. papaya*. *Horovitzia, Jarilla* and *Jacaratia* are the closest genera to *Carica*, and are native to southern Mexico and Central America. *Vasconcellea* is the most closely related genus in South America (52). Twenty-two viral species are reported to cause disease in papaya, and papaya was the host from which eleven of these were first reported (8, 16, 17, 53, 54). For example, PMeV has only been reported to infect papaya (16, 17). Recently, PMeV-2, isolate PMeV-MX, identified in papaya in southeastern Mexico, was reported to also infect watermelon (18). PapMV, in addition to infecting papaya, has been reported naturally infecting pumpkin (*Cucurbita pepo*), *Cnidoscolus chayamansa*, and *Jacaratia mexicana* (10, 11). Mixed infections of PRSV and PapMV have been reported in *C. moschata*, *C. pepo* var. *cylindrica*, and *Citrullus lanatus* (11). Two strains of PRSV (P and W) have been reported, distinguished by host range, where PRSV strain P can infect papaya and cucurbits. In this analysis, we recovered PRSV only from papaya plants (in both regions), with the exception of recovery from insects in the Central Depression.

We recovered four novel viruses infecting papaya and two new reports for papaya. We report two novel begomoviruses provisionally named PapBV 1 and 2, recovered from papaya plants in both regions. Both sequences shared homology to segment A of the genus *Begomovirus*, with lengths 2.7 and 2.8 kb. Two viruses, LCV and EuMV, are new reports for papaya in Chiapas. The crinivirus, LCV, and a begomovirus, TYLCV, were recently reported in papaya in Southern Texas, causing severe symptoms in papaya (8). Additionally, we identified a novel nucleorhabdovirus by TEM images with pleomorphic virions (Fig. S1), and sequences generated by HTS, up to 5.4 kb, isolated from papaya in the Pacific Coastal Plain. Only one rhabdovirus has been reported in papaya, causing apical necrosis disease, from Florida and Venezuela in the 1980’s (12, 55). In the late 1990’s, similar symptoms were reported in southeast Mexico (15), however, there are no rhabdovirus sequence accessions submitted from papaya. We suggest that PNRV-1 identified in Chiapas could be the causal agent of the papaya apical necrosis, although further information is required to confirm this hypothesis. Additionally, in this study, we report novel viruses sharing characteristics with bipartite comoviruses, provisionally named papaya comovirus 1 and 2 (PCV-1, Acc. No. RNA1, pending and RNA 2 pending), in papaya and weeds in the Pacific Coastal Plain. Interestingly, LCV, EuMV, PNRV-1 and PCV 1 and 2, together with PapMV and PMeV, were associated with PRSV.

Viruses may be protective agents, where interactions between viruses within the host are important determinants of the severity of infection, and a wide range of interaction types are possible (21, 56–59). Notice the similar composition of RNA and DNA viruses in the papayas with symptoms of PRSV in both the Pacific Coastal Plain and the Central Depression. Interestingly, PapMV was only reported in papaya plants with non-PRSV symptoms in the Central Depression, however, the role of PapMV as a protective agent was not evaluated *in situ*. We have previously reported changes in the symptomology with either a co-inoculation or a stepwise inoculation, where PRSV followed by PapMV caused synergism, and the reciprocal stepwise inoculation of PapMV followed by PRSV led to antagonism (20). This observation suggested that PapMV could interact as a protective agent against PRSV, and probably against other viruses, as well. PapMV triggers systemic acquired resistance (SAR) in papaya, increasing the expression of a marker protein related to pathogenesis (PR1), which inhibits subsequent infections by PRSV (20).

PapMV-triggered SAR may be a mechanism of plant defense against PRSV, if papaya lacks a set genes conferring resistance to ringspot disease (20, 52). Future studies of the virome at the individual host level, as compared to bulked samples, will help to clarify virus interactions. It would also be interesting to study these interactions at the molecular level, to identify antagonistic or synergistic effects in individual papaya plants, as recently reported for PapMV interactions (20).

### The papaya virome network: metrics for disease management

Bipartite network analysis has been used to characterize complex systems of trophic networks, such as plants and pollinators (including bees, bats, and birds), hosts and parasitoids (including fish and mammals, and ecto- and endoparasites), and bacteria and bacteriophages (29, 30, 60–62). We developed applications of bipartite network analysis for plant obligate intracellular parasites, for the example of viruses in interactions with plants and vectors. This bipartite network analysis includes both illustration of the network structure in images, and quantitative analysis of the network structure. Plant viral metagenomics techniques are sensitive enough to reveal most viral sequences within a plant (23, 63). Also, interpretation of these metrics needs to take into account sampling effort, plant abundance, and insect behavior, such as the actions of vector species (64).

Bipartite network analysis of the papaya orchard virome also focused on host-locations, and a one-mode projection of the network was generated. Betweenness centrality measures the role of a node as a bridge for paths through the network from one host-location to another host-location. Information about the role of hosts and viruses in the network can be translated into potential risk management strategies. For example, weeds and visually asymptomatic papayas in the Pacific Coastal Plain had the highest betweenness centrality (Table 2), however, the highest number of shared viruses within the Pacific Coastal Plain was for visually asymptomatic papaya and papaya with PRSV symptoms (Fig. 6, blue nodes). This suggests that asymptomatic papaya plants and plants with PRSV symptoms are both contributing as a source of viruses.

**Fig. 6.**
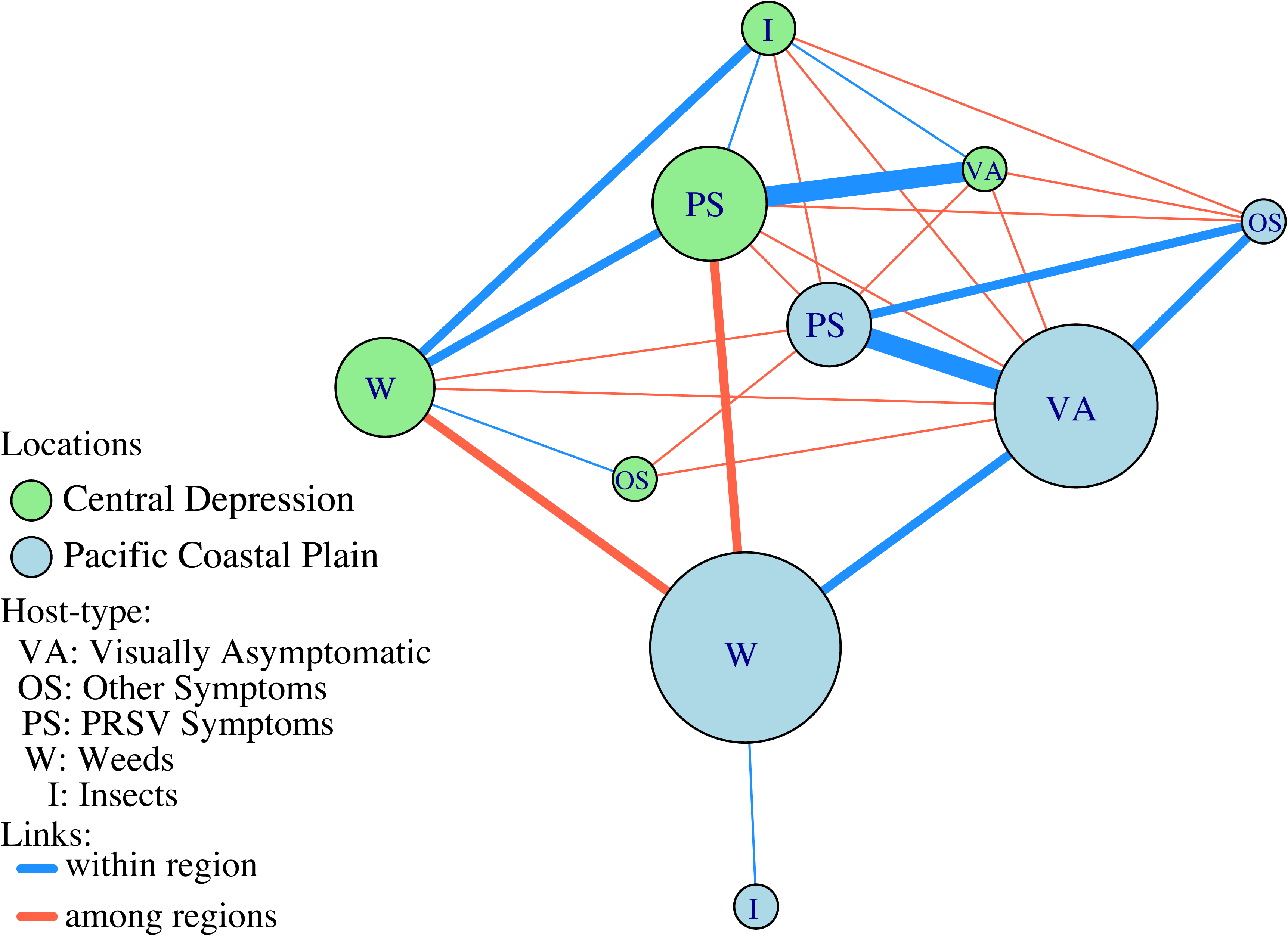
A one-mode network extracted from the bipartite network of associations. Nodes represent host types from the Central Depression and the Pacific Coastal Plain. Node size is proportional to the betweenness centrality of the node. Links represent associations of viruses, and link thickness is proportional to the number of viruses shared within (blue) and among (red) regions.

A scenario consistent with our expectations was observed in the Central Depression, where papaya with PRSV symptoms had the highest betweenness centrality, followed by weeds and insects. Asymptomatic papaya and non-PRSV symptomatic papaya had betweenness centrality equal to 0 (Fig. 6, green nodes). These results suggest that management strategies may be different for each region. The Pacific Coastal Plain is a more fragmented ecosystem, and heavily managed, compared to the Central Depression. Other agroecological differences between the two regions, such as farm size, diversity of plants, and management strategies, may also play a role in the virome dynamics in the papaya orchards. Translation of these results could include future studies to evaluate the cost-effectiveness of weed management, and to consider diagnostic assays to evaluate viral thresholds in visually asymptomatic papayas.

### Virome perspective

The phytobiome is conceptualized as the interactions among microorganisms, the environment, and plants (65). Virome interactions in marine ecosystems have been described, but there is limited information about plant pathogenic virus interactions beyond studies of virus pairs. Plant–virus interactions can produce a number of outcomes, and the possibilities increase when there are multiple viruses. Little is known about the virus community in agroecological landscapes, and how the total number of interactions impact the susceptibility of infected plants to other viruses, where initial infection by one species may facilitate the establishment of another virus species (57). Bipartite network metrics help in identifying the properties of the community, including specialization patterns.

Plant-pollinator networks are asymmetric, relying on generalist species to maintain the nested structure of the network that supports specialized species (30, 66). Host–parasitoid networks are often asymmetric as well, with the nestedness of the network having the opposite effect-specialist species tend to parasitize hosts with more parasites, and generalist parasites tend to parasitize hosts with fewer parasites (31, 61). The papaya orchard virome was asymmetric, and slightly modular and strongly nested. Papaya viruses tend to be in a module with papayas. Viruses in weeds were identified in papayas, as well as suggesting the potential for emerging pathogens.

In our study, there was strong evidence for the observed modularity of the papaya orchard virome for null model 1, but not for null models 2 and 3. There was strong evidence for nestedness of the papaya orchard virome for all three null models, indicating that the observed nestedness may be due to first order properties (29). It is important to keep in mind that networks of specialization may differ by region in temperate or tropical areas, affecting the viral community structure (64). It will be interesting to compare the ecological properties, such as nestedness, of the papaya orchard virome network in Chiapas to networks in other papaya orchards, and other global cropping systems. The identification of these ecological patterns in agroecosystems will support understanding of the contrasting dynamics of tropical and temperate systems over time. Future work emphasizing the ecological properties of plant virome networks will support risk assessment for disease emergence, and the discovery and management of new viruses and new virus interactions.

## Materials and Methods

### Sample collection

We collected samples in two physiographic regions in southern Mexico with significant papaya production, separated by a mountain range: the Pacific Coastal Plain, at sea level, and the Central Depression, at 700 masl. The regions have different levels of ecosystem fragmentation, with patches of deciduous forest, secondary vegetation, and farmland in the Central Depression, contrasting with former deciduous tropical forest replaced with grasslands and farms in the Pacific Coastal Plain (67, 68). The samples consisted of leaf pieces of papaya and weeds, and insects, all collected in September 2014 from three papaya orchards in each physiographic region (Fig. 1). The farms were: El Rocio, Ejido Aquiles Serdan and Santa Lucia in the counties of Acapetahua, Mazatán and Suchiate in the Pacific Coastal Plain, and San Juan de Acala, La Unión, Monte Achiote and La Fortuna in the counties of Acalá, Villa Corzo and La Concordia in the Central Depression. At each farm, a zig zag sampling method was used and farmers provided approximately seven young papaya leaves from the top of trees, categorized by the farmers into each of three symptom types: visually asymptomatic, symptoms that are PRSV-like, and symptoms that are non-PRSV. Two samples from each leaf, about 6 cm^2^, were kept in a plastic bag (see below). Weeds were sampled by collecting asymptomatic and symptomatic leaves in the orchards or their surroundings and insect were actively sampled using insect sweep collecting nets by walking along the perimeter of the orchards and in a cross sections inside the orchard and passively pulling out insects from sticky traps placed in the orchard by the owner. Samples from these orchards were weed species in the Euphorbiaceae, Poaceae, Solanaceae, Convolvulaceae, Asteraceae, Lamiaceae, Anacardiaceae, Malvaceae, Salicaceae, Moraceae, Plantaginaceae, Cucurbitaceae, Portulacaceae, Caryophyllaceae and Amaranthaceae families (Table S1), and insects in the orders Coleoptera, Dermaptera, Hemiptera, Homoptera, Hymenoptera, Odonata and Orthoptera (Table S2). Within each region, papaya samples were pooled by each of the three symptom types, weeds were pooled, and insects were pooled. Plant samples were excised with a knife treated with quaternary ammonium salts before each collection. All individual sampling bags were transported on ice. Samples were rinsed with nuclease-free water and stored at −80°C until processing.

### Purification of viral-like particles

To obtain enriched viral nucleic acids, VLPs and double-stranded RNA from plant and insects were extracted. To obtain the VLPs, a mixture of 100 grams of frozen plant tissue from about 14.5 g of each of the seven leaf pieces collected per sample were pulverized and then homogenized with 40 ml of PBS buffer and further addition of 60 µL of 0.25mM of iodoacetamide and 125 µL of Triton X-100 (33%). The homogenate was stirred for 10 minutes and centrifuged at 13,000 xg for 30 min. The supernatant was filtered through a 0.22 µm pore-size sterile filter (Millipore, Billerica, MA) to eliminate particles of higher density and mass including bacteria, eukaryotic cells, or their fragments. Afterwards, VLPs were precipitated with 10% w/v polyethylene glycol 8000 (PEG), an overnight incubation at 4°C, and a further centrifugation at 13,000 x g for 1 h. The pellet was resuspended in PBS buffer and incubated for 15 min with 0.7 v/v of chloroform to lyse any remaining cellular contamination. Insect VLPs were partially purified only with SM Buffer (50 mM Tris-HCl, 10 mM MgSO_4_, 0.1 M NaCl, pH 7.5) (69). Samples of the VLPs were deposited on formvar-coated 200-mesh copper grids and negatively stained in 1% phosphotungstic acid for 10 min. Finally, they were examined by transmission electron microscopy (TEM). Pleomorphic, filamentous and icosahedral particles were visualized (Fig. S1). Once the VLPs were confirmed, we proceeded with the nucleic acids extraction.

### Nucleic acid extraction

Nucleic acids were isolated from the VLPs pellets using a procedure described for the extraction of PMeV-RNA in latex (70). VLPs were incubated for 2 hours in the DNAse and RNAse cocktails (Invitrogen, Carlsbad, CA) with the addition of 14 µL of 20 mg/ml Proteinase K (Thermo Scientific, Waltham, MA) and the mixtures incubated at 37°C for 30 min. Nucleic acid suspensions were extracted with a volume of Tris-HCl (pH 7.5) saturated phenol. After centrifugation at 8,000 × g for 4 min at 4°C, the aqueous phases were transferred to clean tubes and a second step of extraction with chloroform:isoamyl alcohol (24:1) was repeated. One volume of 3 M sodium acetate (pH 5.2), and 2.5 volumes of ethanol were added to the samples. After centrifugation at 12,000 × g for 20 min at 4°C, the pellet was resuspended in 35 µL of RNase-free water.

### Enrichment of dsRNA

The dsRNA enrichment procedure is a microscale adaptation of a published method (71). 5 g of plant tissue per sample mix were flash frozen in liquid nitrogen, pulverized with a mortar and pestle and deposited in a 1.5 ml tube for the immediate addition of 4 v/w of extraction buffer STE (0.1 M NaCl; 50 mM Tris pH8; 1 mM EDTA; 1% SDS), and final addition of 0.1% 2-mercaptoethanol; 1% bentonite and 2 volume of TE saturated phenol:chloroform (1:1), and shaken vigorously for 10 min. The resulting slurry was centrifuged at 8,000 xg for 15 min at 4°C and the aqueous phase was recovered and deposited into a new 1.5 ml tube containing 0.02 g of CF-11 cellulose (Cole-Parmer Scientific, Vernon Hills, IL). Ethanol was added to a final concentration of 16%, the mixture was shaken, and then centrifuged at 8,000 xg for 5 min at 4°C. The cellulose was resuspended thoroughly and the centrifugation process was repeated several times until there were no traces of color. Thereafter, 200 µL of STE were added, and the mixtures were centrifuged at 8,000 xg for 5 min at 4°C. This step was repeated three times, followed by ethanol precipitation of dsRNA at −20°C overnight. After a centrifugation of 10,000 xg at 4°C for 30 min, the dsRNA pellet was dissolved in a volume of 30 µL of RNase-free water and stored at −20°C until further downstream applications.

### First and Second Strand Synthesis from RNA or dsRNA

For the RNA viruses, the first DNA strand was synthesized with the murine reverse transcriptase (Invitrogen, Carlsbad, CA), following the manufacturer’s recommendations for cDNA synthesis with 2 µL of 10 µM dT_18_ primer, or (20 µM) of each of three random primers based on the reported 5’CCTTCGGATCCTCCN_6-12_3’ (63): 5’CCTTCGGATCCTCCGTACTA3’, 5’CCTTCGGATCCTCCGTCTCCATGTAC3’ and 5’CCTTCGGATCCTCCTCTAGT3’. For the dsRNA viruses, 1 µL of dsRNA and 4 µL of viral nucleic acids were combined with 2 µL of those random primers, 5 µL of dNTPs (10 mM each) in a microtube, were denatured at 65°C for 2 min, and subsequently quenched on ice. Then, 4 µL of 5 X First-Strand Buffer (250 mM Tris-HCl, pH 8.3, 75 mM KCl; 15 mM MgCl_2_), 2 µL dithiothreitol and 1 µL of Superscript II (Invitrogen, Carlsbad, CA) were added to each microtube, and then incubated first at 42°C for 60 min and then at 70°C for 15 min. This was followed by alcoholic precipitation using 3M ammonium acetate and absolute ethanol. The second strand cDNA was synthesized according to the manufacturer’s recommendations, 20 µL of purified single stranded cDNA was mixed with 20 µL NEB 10 Xbuffer, 6 µL dNTPs (10 mM), 2µL RNAse H (2U/µL), 3 µL DNA polymerase I (50,000 U/ml NEB) and 80 µL deionized H_2_O. The sample was incubated for 2.5 h at 16°C and purified with phenol:chloroform:isoamyl alcohol (24:25:1).

### Library preparation, sequencing and sequence analysis

Ten samples in total, five sample types from each region, Pacific Coastal Plain, and Central Depression. Three types of papaya symptom types were sampled: visually asymptomatic papaya, papaya with PRSV symptoms, and papaya with non-PRSV symptoms, along with bulked samples of weeds and insects. The duplicated cDNA libraries for each of the ten host-location combinations were obtained using the Nextera XT library preparation kit and sequenced by Illumina HiSeq 2500 fast mode, paired-end reads (2×100) (Cinvestav Sequencing Facility, Irapuato, MX). The total number of raw reads we obtained was 66,406,113, and an average of 2 million reads per library was obtained after removing low-quality reads. The libraries from papaya were filtered using the draft genome sequence of the *C. papaya* reference genome [Acc. No. ABIM01] with BOWTIE2 (72, 73). All libraries were *de novo* assembled using Spades v.3.7 (74) enabling the metagenomics option, yielding a total of 220,792 contigs, where all sequences lower than 500 nt were discarded. The total number of contigs was analyzed with a first iteration of BLASTx (75) using a local database of viruses VirDB (76), where sequences > 95% identity, sequences with homology and e-values > e^-5^ to plant virus were retained. The recovered contigs were searched with a second iteration of BLASTn against the non-redundant database from NCBI with e-values > e^-5^. Sequences belonging to taxa other than plant viruses were discarded. The contigs were linked to scaffolds with Geneious v.R11. Contigs and scaffolds recovered were used for re-aligning the paired-end reads using a custom pipeline implementing trinity to estimate the abundance of reads, and calculating the stats with bbmap (sourceforge.net/projects/bbmap), using BOWTIE2, and samtools (77). All sequences generated for downstream analysis were nearly full length (with variations of ±0.2 kb), considering as complete ORFs for RNA viruses, and for DNA viruses two ORFs for geminiviruses, and one ORF for the alphasatellite. The only partial sequence included in the analysis from plants was the rhabdoviruses in papaya, it was visualized by TEM, and validated by RT-PCR.

### Taxonomic analysis

The results of the search for homology by BLAST to known viruses against databases of nucleotide or amino acid sequences allowed the identification of sequences to distantly related viruses with low similarity to higher taxonomic levels (e.g., family or genus), or for sequences with high similarity to known virus species. The sequences assembled were aligned among them using Needleman and Wunsch in Geneious (78). Sequences grouped by similarity were classified taxonomically according to their species demarcation criteria following ICTV rules (https://talk.ictvonline.org/), either by identities using BLAST, or pairwise similarities. Sequences were submitted to NCBI (Table S3).

### Bipartite network analysis

We constructed a bipartite adjacency matrix, where one group was viral taxa and the other group was the combination of host type and region. The entries in the adjacency matrix indicated the presence or absence of each viral taxon in each host-location combination. We also considered weighted matrices with weights representing the read coverage across the contig length as an indicator of relative sequence abundance. Network metrics such as node degree, nestedness, and others were calculated using the bipartite package in R (79). Hypothesis tests for modularity and nestedness of networks were evaluated using a standardized effect size (SES). A Z-score test was calculated for these two metrics as follows: Z = (observed metric – mean under null) / standard deviation under null. Calculations used a two-tailed normal distribution, compared to three null models. Null model 1 maintained sample richness, null model 2 maintained species frequency, (37, 80, 81), and null model 3 maintained both species frequency and sample richness (41). 1000 matrices were generated with 100 iterations each, using the picante package in R (82). Bipartite network figures were generated using the igraph package in R using the Davidson-Harel (DH) algorithm layout (83, 84). Pie plots were generated with sunburstR, and other figures were generated with ggplot, these analyses were conducted in the R package v.3.4.2 (40).

## Acknowledgments

This work was carried out with support from Agromod through sponsorship from a PEI-PROINNOVA program by CONACYT, Mexico. RI. Alcalá-Briseño Ph. D. scholarship (No. 410151) was sponsored by CONACYT. We also appreciate support from the University of Florida. We thank Javier Luévano-Borroel from CINVESTAV-IPN for the identification of the insect families. We thank José C. Hughet-Tapía for helpful discussions about the bioinformatic analysis, and Masatoshi Katabuchi for helpful discussions about the ecological models.

